# Reassessing the Revolution’s Resolutions

**DOI:** 10.1101/224402

**Authors:** Marin van Heel, Michael Schatz

## Abstract

***We are currently facing an avalanche of cryo-EM (cryogenic Electron Microscopy) publications presenting beautiful structures at resolution levels of ~3Å: a true “resolution revolution” [Kühlbrandt, Science 343(2014)1443-1444]. Impressive as these results may be, a fundamental statistical error has persisted in the literature that affects the numerical resolution values for practically all published structures. The error goes back to a misinterpretation of basic statistics and pervades virtually all popular cryo-EM quality metrics***. The resolution in cryo-EM is typically assessed by the Fourier Shell Correlation “FSC” [Harauz & van Heel: *Optik* **73**(1986)146-156] using a fixed threshold value of 0.143 (“FSC0.143”) [Rosenthal, Henderson, *J. Mol. Biol.* **333**(2003)721–745]. Using a simple model experiment we illustrate why this fixed threshold is flawed and we pinpoint the source of the resolution confusion. When two vectors are ***uncorrelated*** the expectation value of their inner-product is zero. That, however, does not imply that each individual inner-product of the vectors is zero (the vectors are ***not orthogonal***). This error was introduced to electron microscopy in [Frank & Al-Ali, *Nature* **256**(1975)376-379] and has since proliferated into virtually all quality and resolution-related metrics in EM. One criterion not affected by this error is the information-based ½-bit FSC threshold [van Heel & Schatz: *J. Struct. Biol.* **151**(2005)250-262].

---

The FSC metric was introduced for comparing the information content of two volumes as function of spatial frequency [Harauz & van Heel 1986]. It is not restricted to cryo-EM or structural biology and can be used to assess the reproducible resolution of results from all instruments or algorithms that produce three-dimensional (3D) information. The FSC metric compares the full data (phases and amplitudes) and does not depend on any *a priori* assumption on the nature of the signal or noise in the volumes. The FSC is a one-dimensional curve showing a high correlation level (~1) at the low resolution end, gradually dropping towards zero at higher spatial frequencies, and finally oscillating around the zero mark close to the Nyquist frequency where only random correlations should be present [van Heel & Schatz 2005]. For extracting a representative *single* resolution value from an experimental FSC curve, a sensible threshold value must be agreed upon. Defining such a threshold, *does* require assumptions to be made about the nature of the noisy 3D volumes being compared.

The standard assumption made here is that the two volumes (Fig 1a-b) contain identically the same information (the signal “**S**”, Fig 1d-e), while each is deteriorated by a different realisation of additive (zero mean) random noise (“**N_i_**”, Fig 1f-g). In our simple test illustrating the basic principles, “**S**” is the structure of the worm hemoglobin [Afanasiev 2017], but that is immaterial to our model experiment. The two volumes: **A = S + N_1_** (Fig 1a), and **B = S + N_2_** (Fig 1b), were generated using volume S and the two random-noise volumes **N_1_** and **N_2_**, respectively. From the 3D Fourier transforms of these 3D densities (transforms denoted in italics), the shell correlations are calculated: 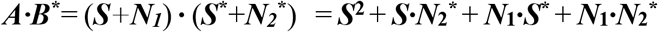. The following FSC curves (function of radius ***r***) are generated:

1) A “conventional” FSC_AB_ (Fig 1c) between the two noisy volumes ***A*** and ***B***,
2) The “signal-versus-noise” correlation: FSC_SN1_ between ***S*** and ***N_1_*** (Fig 1i),
3) The “signal-versus-noise” correlation: FSC_SN2_ between *S* and *N_2_* (Fig 1j),
4) And the noise-versus-noise correlation: FSC_N1N2_ (Fig 1h), between ***N_1_*** and ***N_2_***.

**Figure 1.**
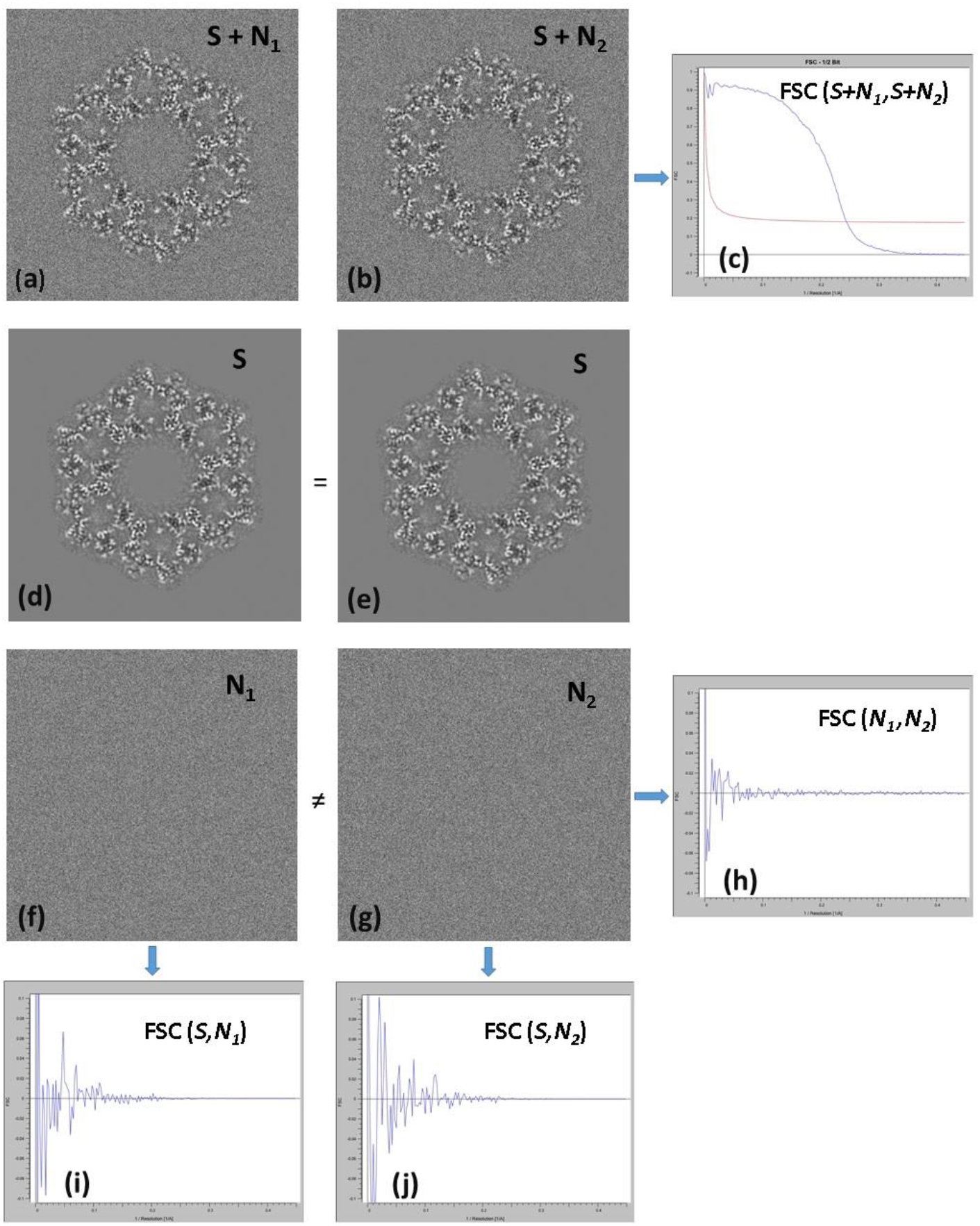
Fourier Shell Correlations between Signal and Noise volumes. **a)** Section through the test volume **A**: the sum of the fixed signal **S** and random noise volume **N_1_**. Only one section (150/360) of the 3D volumes is depicted throughout. **b)** Corresponding section through test volume **B**: sum of signal S and noise **N_2_**. **c)** FSC between the two noisy volumes **A** and **B**. **d)** and **e)** Two identical copies of signal **S** (an arbitrary cryo-EM 3D reconstruction). **f)** Corresponding section through noise volume **N_1_**. **g)** Corresponding section through a second noise volume **N_2_**. **h)** FSC (not normalised) between noise volumes **N_1_** and **N_2_**. **i)** FSC (not normalised) between signal **S** and noise volume **N_1_**. **j)** FSC (not normalised) between signal **S** and noise volume **N_2_**. (Note that Figures: **1h, 1i**, and **1j,** have the same vertical scale). All calculations were performed using the IMAGIC-4D software system [van Heel 2012].

(Note that the FSC_SN1_, FSC_SN2_, and FSC_SN2_ curves are normalized only by the number of voxels in the shells and are therefore directly comparable). The FSCab (Fig. 1c) has a typical FSC appearance indicating that realistic levels of noise have been added to the signal ***S*** to create the noisy volumes ***A*** and ***B***. The noise-versus-noise FSC_N1N2_ (Fig 1h) oscillates around the zero mark over all spatial frequencies with its modulation increasing close to the origin, where Fourier shells contain fewer voxels. Note that the correlation fluctuations between the signal ***S*** and each of the noise terms ***N_1_*** and ***N_2_*** are significantly ***larger*** than are the correlations between ***N_1_*** and ***N_2_***. Whenever the signal ***S*** and the uncorrelated noise ***N_i_*** are of a similar standard deviation, the predominant noise component consists of the cross-terms 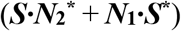 rather than of only the smaller 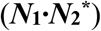 correlations. Resolution criteria must compare the overall signal with the influence of *all three* of these noise-related cross-terms.

It is at this level of the theoretical considerations that a fundamental statistical error is made in the literature. It is assumed that since the signal and the noise are ***uncorrelated***, their inner products 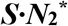 and ***N_1_·S\**** are zero, and the noise-versus-signal cross-terms can subsequently be dropped from all equations. However, “***uncorrelated***” means that the inner product between these vectors is zero as an ***expectation value***, that is, as the average over an infinite number of repeats of the experiment (symbolically: <***S·N***> = **0**). Assuming that ***each*** of the “***S***·***N***” inner products is zero is a very different matter; it implies that ***each*** of these vector pairs are ***orthogonal***. The latter assumption – prolific in the EM literature – is untenable as is clear from the model experiment. Thus, although the statement that the ***S***ignal and the *N*oise are ***uncorrelated*** is by itself correct, the interpretation of its practical consequence, that each of the Signal and the Noise vector-pairs are ***orthogonal***, is fundamentally flawed.

This assumed orthogonality is implicit in virtually all papers on FSC resolution thresholds, SNR ratios, Spectral SNRs, etc. This includes all papers where fixed-valued FSC resolution thresholds are proposed. The primary resolution-determining noise factors are the ***S·N*** cross-terms which hitherto have been left out of the equation by a semantic misinterpretation of the underlying statistics, including the work by Rosenthal & Henderson [2003], which provides the basis for the popular **0.143** FSC threshold. This statistical error was introduced to the EM literature by Frank & Al-Ali [1975] and has since proliferated in the field. Frank & Al-Ali [1975] related the signal-to-noise ratio SNR (“**α**”) to the cross-correlation coefficients (“***ρ***”), and the resulting formula (***α=ρ/(1-ρ***)) is normally used to justify resolution-related metrics (including spectral SNRs [Baxter 2009]). A basic problem with this formula is readily seen by substituting a “-1” for the value of ***ρ*** (range: −1 ≤ ***ρ*** ≤ +1), leading to an SNR value of −0.5, impossible for a positive entity. Negative SNRs result for all negative ***ρ*** values; these occur frequently, especially when the signal is small.

To pinpoint the exact mistake made by Frank & Al-Ali [1975] requires a look at the underlying mathematics, which were copied from an earlier paper [Bershad & Rockmore 1974]. This original paper was based on very strict (yet unrealistic) *a priori* orthogonality assumptions, meaning that – mathematically correct or not – it cannot be used in practice. Their orthogonality assumptions are in contradiction with their other assumption that the signal and noise are band-limited *(Supplementary Materials).* Frank & Al-Ali applied the formulas to practical EM data, thus violating the orthogonality assumptions. The justification given for extending this applicability was: “In image processing application where N is of the order of 10,000, the Gaussian assumption can therefore be dropped” *(Supplementary Materials).* Saxton, in his [1978] book, provides a comprehensive derivation of the (Bershad & Rockmore/Frank & Al-Ali) formulas illustrating when what cross-terms are assumed zero. In all these papers, however, the same mistake is made explicitly or implicitly, namely: concluding from a ***zero expectation value*** of the cross-terms, that ***each*** of the measured cross-correlation terms is ***zero***, thus ignoring the main noise source in the analysis.

We have criticized these errors previously in [van Heel & Schatz 2005], but our findings have largely been ignored (see for example: [Liao & Frank 2010; Scheres & Chen 2012, Chen *et al.* 2013]). (Liao & Frank [2010], do cite our [2005] publication, but not for its actual scientific content). We emphasize that the popular fixed FSC threshold criterion of “0.143” of Rosenthal & Henderson [2003] is itself incorrect – even in an asymptotic sense – because the ***S·N*** cross-terms have explicitly been left out during its derivation. The power in the ***S·N*** cross-terms is determined by the standard deviation of signal ***S*** multiplied by the standard deviation of noise ***N*** [van Heel & Schatz 2005] as a direct consequence of the central limits theorem [Feller 1945; Wikipedia 2017]. The region in Fourier space that is important for determining whether one has already collected enough signal ***S*** from the noise background ***N*** is just that region where the ***S·N*** cross-terms represent the main source of noise variance. This primary noise source has been ignored in the fixed-value resolution thresholds. The 1/2 -bit FSC criterion [van Heel & Schatz 2005] *does* include the influence ***S·N*** cross-terms correctly and is not dependent on any comparison to historical phase-oriented metrics from X-ray crystallography. It is currently the only FSC threshold with a justified asymptotic behaviour, asymptotically approaching the numerical value of **0.1716**.

The statistical error affecting resolution metrics in the cryo-EM literature persists in a second generation of metrics that builds upon these metrics. The FSC, for example, has been used to assess ***local*** resolutions in 3D maps [Cardone 2013]. Using a fixed threshold value (FSC0.5 in this case), as promoted by [Rosenthal & Henderson 2003], implies that *different* areas of the 3D reconstructions are essentially assessed with *different* resolution criteria *(Supplementary Materials).* Such programs will necessarily yield inconsistent results. Similarly, when the FSC is used to assess the effects of preferential orientation, a correct threshold curve must be used instead of a fixed-value FSC threshold [Afanasyev 2017]. New FSC-based structural refinement approaches [Murshudov 2016] are also hampered by those erroneous statistics.

The fundamental problems that cryo-EM has slipped into can be seen as a semantic confusion of the underlying mathematics: ***uncorrelated*** measurements are not necessarily ***orthogonal***. New confusions have been added recently to the cryo-EM resolution discussion. The “gold standard FSC” [Scheres & Chen 2012] may be taken as an implicit claim of superiority for their specific FSC metric, since “*gold standard*” is used as an adjective to “FSC”. However, this *“goldstandard*” reflects the cryo-EM-specific alignment procedures, and the “FSC” referred to is identically the same *“old standard*” FSC [Harauz & van Heel 1986], albeit used with an incorrect 0.143 threshold. A more extensive list of papers directly affected by the Frank & Al-Ali [1975] error can be found in the *Supplementary Materials.*

The Rosenthal & Henderson [2003] fixed-threshold criterion is intimately linked to the Figure of Merit (FOM) used in X-ray crystallography. Structural biology has been dominated by X-ray crystallography for decades, and in X-ray crystallography the primary data are ***amplitudes***, and to which matching ***phases*** have to be found. The quality metrics in X-ray crystallography have thus traditionally focused on that phase determination. The popular R-factor measures the difference between the observed amplitudes and the amplitudes calculated from an atomic model which, in turn, is primarily determined by its phases. The R-factor is a rather indirect, crystallography-specific metric that includes the iterative fitting of atomic co-ordinates of (biological) macromolecules into an electron density that, in turn, is iteratively modified to accommodate that atomic model. The FOM is another such phase-oriented, crystallography-specific metric. The early quality criteria in EM naturally followed their crystallography “big brother” examples, using phase residuals and the highest visible 2D crystal diffraction peaks as resolution criteria.

The fact that EM produces images (and not just diffraction patterns) has the great advantage that both amplitudes and phases can simultaneously be used in its quality metrics. The FSC does just that and is therefore a *better* metric than are any of its X-ray crystallography predecessors [van Heel 2000]. In spite of electron microscopy (EM) historically producing lower resolution results than X-ray crystallography, the EM FSC metric is a much more reliable general-purpose resolution yardstick than the traditional crystallography metrics. One can have poor metrics defining the quality of a beautiful X-ray map at ~2Å-resolution and at the same time have an excellent metric defining the quality of a mediocre ~20Å resolution cryo-EM map [van Heel & Schatz 2005]. It is only now that X-ray crystallography and cryo-EM are yielding structures at comparable resolution levels that the quality of the resolution *metrics* in both fields are being discussed again. Trying to curtail the richer FSC metric to match a phase-oriented X-ray crystallography metric (FOM), in order to justify a fixed-value threshold for the FSC [Rosenthal & Henderson 2003] is an anachronism, even if no statistical errors had been made in the process.

The FSC has the great advantage that it is a universal metric applicable to all experiments that generate 3D data, without specific ties – other than historic ones – to structural biology. It is rewarding to see that FSCs and FRCs (the 2D version of the FSC) are now increasingly making their way into other fields, including X-ray crystallography, X-ray tomography, super resolution light microscopy, and even planetary sciences [Afanasyev 2015; Murshudov 2017, Vila-Comamala 2011].

In most EM resolution-metric papers, the fundamental statistical error of (implicitly or explicitly) assuming orthogonality between signal and noise is made in the first paragraphs of the paper (or earlier, when building upon previous publications). In some cases, like that of the spectral SNR metric, one needs to think twice to precisely find where the implicit orthogonality assumption has been made *(Supplementary Materials).* This orthogonality assumption invalidates the numerical values of the associated resolution claims, especially when the number of measurements (pixels/voxels) is small as is the case in local resolution estimation. These historical errors are impeding methodology developments and should therefore be overcome by adopting quality measures not affected by these errors.

Our criticism of the assumptions underlying most current resolution metrics does not mean that we are questioning all cryo-EM *biological results:* when reliable side-chain information is available, for example, the exact numerical resolution labels are only of secondary importance. What we are stating is that the popular fixed-value FSC threshold criteria (0.5 or 0.143) are based on incorrect mathematical assumptions and cannot be used to compare independently conducted cryo-EM experiments. Different areas of the same 3D reconstruction in local FSC measurements already count as “independently conducted experiments”. Let us recapitulate that for any correct resolution criterion the threshold must be dependent on the radius in Fourier space; the criterion must include the influence of the relative box size (and of the pointgroup symmetry). The dataset may not have suffered from alignment reference bias, or from imposing sharp identical masks on both 3D maps to be compared. Last but not least, the data must not be under-sampled. The under-sampling rule means that resolution values should not be assessed beyond ~2/3 of the Nyquist frequency where systematic interpolation artefacts may start accumulating [van Heel & Schatz 2005].

## Acknowledgements

We thank Ralf Schmidt for his support with the model calculations. We thank various colleagues, ex-students and post-docs for discussions on these matters over the last decades, some of whom have explicitly commented on our manuscript. (Names withheld for confidentiality).

## Supplementary Materials

### Introduction

Structural biology & has for decades been dominated by X-ray crystallography. The cryo-EM community was originally not taken too seriously and its results were originally described as “blobology” by the structural biology community. The quality criteria in EM were therefore also not taken very seriously compared to the established X-ray crystallography metrics. In the shadow of the prevalent X-ray crystallography approach, a large number of different quality criteria emerged in EM that crystallised into the final predominance of the FRC and FSC. In covering these developments here, we have used “Google Scholar” to search for the citations in the literature. Those citations also allows us to trace the more recent developments affected by the earlier methodology flaws. The list of relevant papers below is necessarily incomplete and includes only the cryo-EM literature. We invite the readers to trace the citation indices for finding further research affected.

### How large is the number N?

The number of pixels/voxels in a real space image/volume versus the number of pixels/voxels in their Fourier transforms is an important issue in these matters. According to their own claims, the Frank & Al-Ali [1975] mathematics apply to real space images with a large number of **N** pixels: *“In image processing application where N is of the order of 10,000...”*). This reference (and the formula therein) is typically used to justify the dropping of the cross-terms in the theoretical FSC fixed-threshold discussions (for example [Penczek 1988]). This sounds reasonable because the overall number of **N** pixels/voxels in real space is the same as the number of ***N*** pixels/voxels in Fourier space. However, in the case of the FRC/FSC one considers only the *small* number ***N_r_*** of Fourier-space pixels/voxels that fall within a narrow ring/shell ***r*** of the full 2D/3D dataset. In the FRC case ***N_r_ ~ r·N^1/2^***, and in the FSC case ***N_r_ ~ r^2^***·***N*^2/3^**. Because these numbers ***N_r_*** are so much smaller (and are thus even very much smaller close to the origin in Fourier space) than the corresponding real space numbers **N**, the applicability of the large **N** rule [Frank & Al-Ali 1975] to the FSC/FRC cannot not be justified in general, even if the “**α=*ρ***/(***1-ρ***)” formula of Frank & Al-Ali had been correct. This problem is exacerbated when in the estimation of local resolution where the considered volumes are consists of only a small fraction of the original reconstruction.

### Discussions in mailing lists

We wish to point to various discussion threads by different authors in the 3DEM and CCPEM mailing list communities:

3DEM: (<u>http://mail.ncmir.ucsd.edu/pipermail/3dem</u>)

CCPEM: (<u>https://www.jiscmail.ac.uk/cgi-bin/webadmin?A1=ind1705&L=CCPEM</u>), that have touched upon the resolution issues discussed in our paper. Those threads include: August 2013 (“Don’t blame your thermometer…”),

August 2015 (“On FSC curve (A can of worms…)”),

May/June 2017 (“Minimum standards for FSC reporting?”),

June 2017 (“Instrumental Resolution versus Results Resolution”).

Most of these discussion threads can now be closed.

### Details of the various FSCs

In Figure 1 of the main paper, the “normal FSC” is normalised whereas all others are not. The FSC ½-bit threshold is only relevant in the normalized “conventional” (***S+N_1_***) *versus (**S+N_2_**)* FSC. For comparison, we here show the various FSCs separately to illustrate the strong increase in the *(**S·N**)* cross-terms at resolution levels around the ½-bit threshold. At frequencies well beyond the ½-bit threshold the noise-to-noise correlation is the larger noise-contributor. Around the ½-bit threshold the signal-versus-noise cross-terms become the largest contributors to the three overall noise terms.

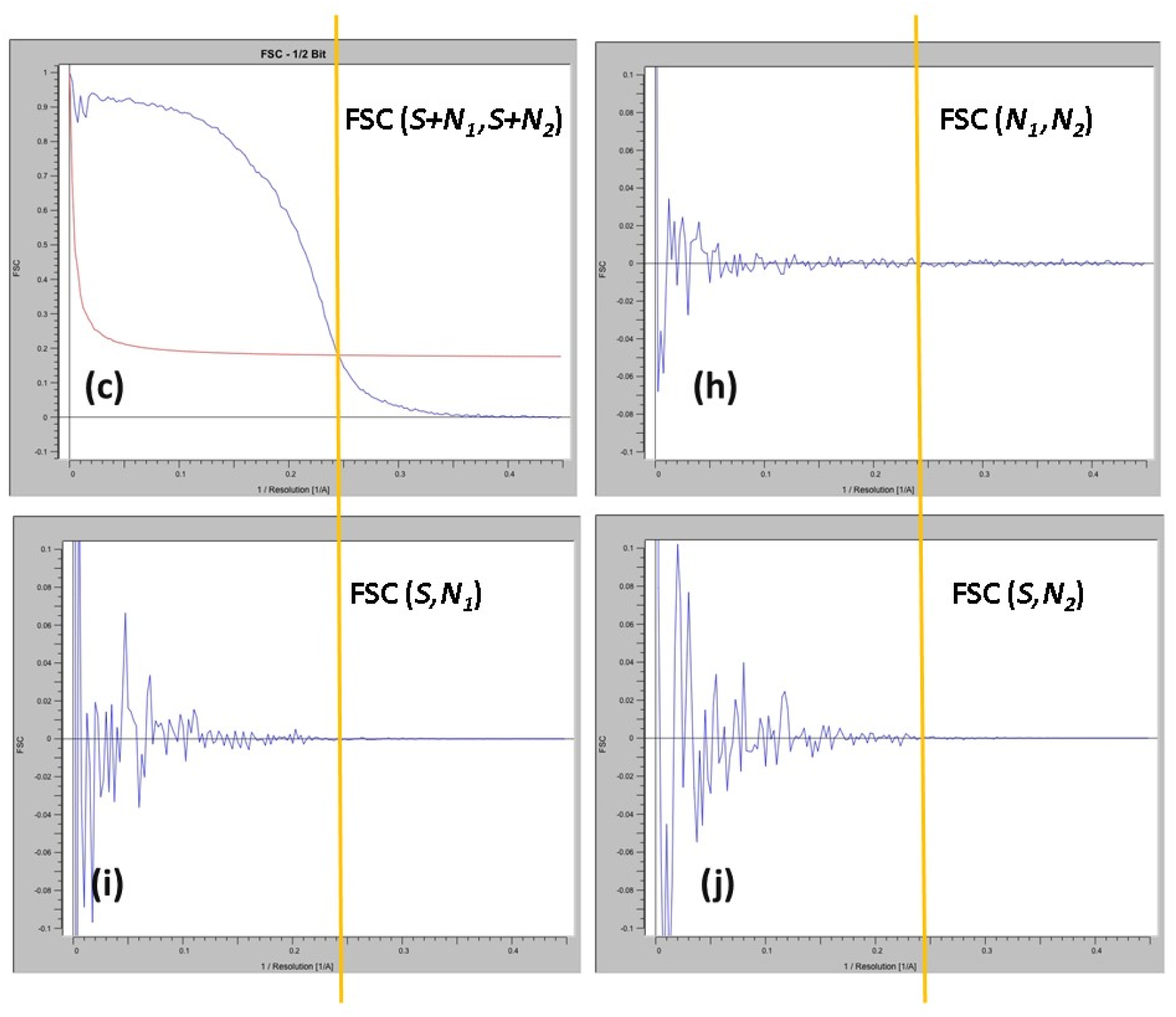

## Cryo-EM resolution metrics literature (Chronological order by year)

Bershad NJ, Rockmore AJ: **On estimating signal-to-noise ratio using the sample correlation coefficient**, *IEEE Trans. Inf. Theory* **20** (1974) 112–113.

> An interesting formula is derived relating the signal-to-noise ratio (SNR) of data to the cross-correlation coefficient between two independent measurements. However, very strict orthogonality properties for the signal and the noise components of the data are postulated. The first assumption is that both the time domain signal ***S***(t) and the noise ***n***(t) are *band limited* (bandwidth **B**) and are sampled at **2B** or better following the rules of the sampling theorem. Two measurements are made in parallel: ***x***(t) = ***s***(t) + ***n_1_***(t), and: ***y***(t) = ***s***(t) + ***n_2_***(t), and those are correlated with each other to come to an estimate of the SNR. Further assumptions made are that, for two measurements not initiated at the same time, the correlation is zero: <***x_i_x_j_***> = δ_ij_; <***y_i_y_j_***> = δ_ij_ ; and <***x_i_y_j_***> = ρδ_ij_, with δ_ij_= 0 for all **i**≠**j**. Let us then focus on the case of ***j=i+1***, that is, the next sample in the time series. Because ***x(t)*** is band-limited and sampled at ***2B***, the signal is zero at the Nyquist frequency: subsequent time samples are therefore correlated and the <***x_i_x_i+1_***> type terms will necessarily be non-zero. The *a priori* assumptions in this paper thus contradict each other, rendering the theory not applicable in practice. With these strict *a priori* assumptions, the issue of the cross-terms between signal versus noise, and the fact that one cannot apply the expectation values to the individual measurements, never came to bear. The resulting formula ***α= ρ/(1-ρ)*** is incorrect, as is discussed below.

Frank J, Al-Ali L: **Signal-to-noise ratio of electron micrographs obtained by crosscorrelation**. *Nature* **256** (1975) 376-379.

> By applying the Bershad & Rockmore [1974] formulas to real experimental electron microscopical data – and explicitly claiming a much larger field of application – these authors violate the mathematical *a priori* assumptions of the original paper. The formula relating cross correlation coefficient ***ρ*** to the SNR (Signal to Noise Ratio) **α** in this paper is: ***α= ρ/(1-ρ)***. Note that **α**, the SNR (S^2^/N^2^) is *per definition* a positive entity. The ***ρ*** in contrast, is defined in the range -1 ≤ ***ρ*** ≥ +1 from full anticorrelation to total correlation. Let us assume that in one case the two measurements are indeed fully anti-correlated, i.e.: ***ρ***=-1. The Frank & Al-Ali [1975] formula then yields: **α** = -1/2, in direct violation of the SNR’s positivity. The problem occurs for any (even very small) negative value of the cross-correlation ***ρ***, a routine occurrence for small signals within large noise fluctuations. Of course, since the formulas in this paper were taken directly from Bershad & Rockmore [1974], this problem was a preexisting condition for which these authors could be excused, had they not incorrectly extended the validity range of the formula to real world experiments. Moreover, we could not find any citations to the original [Bershad & Rockmore 1974] paper from this group after 1980 (only to [Frank & Al-Ali 1975], as in: [Frank 1998; Baxter 2009]) thus assuming ownership of the “**α**=***ρ/(1-ρ)***” formula.

Saxton WO: **Computer Techniques for Image Processing in Electron Microscopy**. Advances in Electronics and Electron Physics (Supplement 10) (1978) Academic Press, New York. Pages 203-205.

> Saxton re-derives the Bershad & Rockmore [1974] and the Frank & Al-Ali [1975] formulas from scratch. Whereas Frank & Al-Ali have only copied the Bershad & Rockmore formulas and verbally extended its validity range, Saxton’s real-space derivation is a transparent treatment of all stages of the background statistics. There is here no explicit mentioning of “expectation values” only of “spatial averages”. Thanks to this clean discussion it becomes very clear when and what cross-terms are assumed to be zero. As is argued in our main paper, those cross-terms (equations 9.4.8 and 9.4.18-9.4.19) should not have been removed because the **S·N** cross-terms are actually the main source of noise-variance in the data.

Frank J, Verschoor A, Boublik M: **Computer averaging of electron micrographs of 40S ribosomal subunits**. *Science* **214** (1981) 1353-1355.

> Introduction of the Differential Phase Residual (DPR) for comparing two 2D images (or averages) with each other as function of spatial frequency. This DPR definition included sums of Fourier amplitudes, making the results dependent on the relative scaling of the densities in the two images, and invalidating the DPR as a general-purpose resolution criterion. This resolution metric was criticized implicitly by the introduction of the FRC [van Heel 1982] and the SFCF [Saxton & Baumeister 1982], and explicitly in (Orlova et al., 1997; Van Heel, 1987).

Van Heel M, Keegstra W, Schutter W, van Bruggen EJF: **Arthropod hemocyanin studied by image analysis**. In: **The Structure and Function of Invertebrate Respiratory Proteins**. (Editor: Wood EJ), EMBO Workshop, Leeds, *Life Chemistry Reports*, *Suppl.* 1, (1982) 69-73.

> This paper is the first – name defining – reference to the Fourier Ring Correlation FRC. Its first use was to compare the results of 2D EM data processing with the published results of X-ray crystallography. The advent of the 3D version of the FRC (that is: the FSC) is already announced. Saxton & Baumeister [1982] independently introduced the same concept.

Saxton WO, Baumeister W, 1982. **The correlation averaging of a regularly arranged bacterial envelope protein**. *J. Microsc.* **127** (1982) 127–138.

> This paper introduces the “spatial frequency correlation function” which is equivalent to the “Fourier Ring Correlation” (FRC [van Heel 1982]). No formulas are given to define their metric but the idea is described correctly. A new concept is introduced, namely that of the significance threshold of the signal reaching more than two standard deviations above the background noise in a certain frequency band. This 2σ-criterion introduces the concept of a 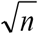 dependency into the resolution criteria in electron microscopy where *n* is the number of pixels within each integration ring.

Harauz G, Van Heel M: **Exact filters for general geometry three dimensional reconstruction**, *Optik* **73** (1986) 146-156.

> This is the first mentioning of the Fourier Shell Correlation (“FSC”) in the literature defined as:
>
>
> 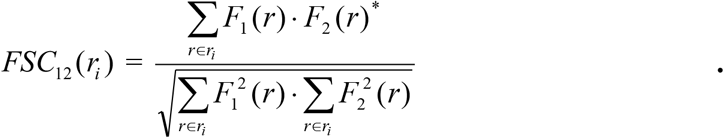
>
> This metric had already been announced as the 3D generalisation of the 2D Fourier Ring Correlation (“FRC”) in [van Heel 1982]. The FSC was here used to assess the quality of 3D reconstruction algorithms, that is, to assess the “instrumental” quality of the algorithm by comparing the original 3D “phantom” with its 3D reconstruction. In such experiments, noise plays no role and it is normally noise that limits the quality of practical 3D reconstructions in cryo-EM, see for example: [van Heel 2000b]. In this first use, the entire FSC curve is used for evaluating the reconstruction quality of an algorithm and the issue of a specific resolution “threshold” is not yet discussed. Full, independently determined data sets were being compared and the cryo-EM-specific “half datasets” did not play a role. For the cryo-EM application of this metric see [van Heel 2000b], and [van Heel & Schatz 2005]

Van Heel M, **Similarity measures between images**, *Ultramicroscopy* **21** (1987) 95-100.

> In this paper an extensive comparison is made between FRC/FSC-type of resolution criteria. This paper explicitly pinpoints the shortcomings of the DPR (Differential Phase Residual) which was normalised by a sum of amplitudes and replaces it by a corrected Fourier Ring Phase Residual (FRPR), normalised by a product of amplitudes. The random noise-limits of the DPR [Frank 1981] is calculated to be 103.9° which was previously assumed to be 90°. Moreover, it is also argued that a fixed 45° phase residual (DPR) threshold makes as little sense as a fixed 0.5 FSC threshold: it is sometimes too low and sometimes too high. It cannot account for all the known systematic factors that influence the comparison of two maps including symmetry, degrees of filling of the 3D map, etc. It is remarkable that such fundamental scientific criticism of the DPR metric is still not taken seriously in some quarters [Liao 2010; Sorzano 2017].

Unser M, Trus BL, Steven AC: **A New Resolution Criterion Based on Spectral Signal-To-Noise Ratios**. *Ultramicroscopy* **23** (1987) 39–51.

> First paper proposing an SSNR (Spectral Signal to Noise Ratio) criterion. Compared to the FSC this method of resolution determination is somewhat problematic because neither the Signal nor the Noise are measurable entities and one thus cannot avoid the discussion of how well the experimental reality is covered by the underlying mathematical model. The paper describes the summing in Fourier Space of a large number of images **X_i_**, in order to come to a reliable estimate of the Signal power and the Noise power.
>
> The images **X_i_** are assumed to contain the same signal **S_i_** and an independent additive noise component **N_i_**: {**X_i_ = S_i_ + N_i_**}. Because of the linearity of the Fourier transform, one can perform the summing of the images in Fourier transforms with: {***X_i_ = S_i_ + N_i_***}. After the summing operation one is left with the complex sum of the (***i=1,n***) images: {Σ***X***= Σ***S*** + Σ***N***}. For calculating the power spectrum from this sum, one needs to calculate ***P*** = {Σ***S*** + Σ***N***}·{Σ***S*** + Σ***N***}* where, again, the cross-term issue between the signal term Σ***S*** and noise term Σ***N***^*^ emerges. As was the case for our practical example in the main paper, the most significant noise terms in this power spectrum will here be the two “Σ***S***·Σ***N***\*” cross-terms. These cross-terms are implicitly assumed to be zero since they are not even explicitly discussed in this (and in other) SSNR papers. Thus, the same cross-term negligence occurs in SSNR deliberations as those discussed for fixed-value FSC threshold calculations.

Penczek P, Grassucci RA, Frank J: The ribosome at improved resolution: new techniques for merging and orientation refinements in 3D cryo-electron microscopy of biological particles. *Ultramicroscopy* **53** (1994) 251-270.

> The DPR [Frank 1981] is here generalised to 3D. No mention of the DPR metric having been refuted a decade earlier. It had been refuted implicitly by the introduction of the FRC in 1982 (and the SFCF 1982), and explicitly by correcting its unfortunate definition in the form of the FRPR [van Heel 1987].

Orlova EV, Dube P, Harris JR, Beckman E, Zemlin F, Markl J, van Heel M: **Structure of Keyhole Limpet Hemocyanin Type 1 (KLH1) at 15Å resolution by electron cryomicroscopy and angular reconstitution**, *J. Mol. Biol.* 271 (1997) 417-437.

> Saxton & Baumeister [1982] introduced the concept of the 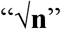 dependency for FSC/FRC resolution type of criteria. That concept is here extended with the idea that the number of independent voxels in a Fourier-space shell reduces with the number of asymmetric units for the given pointgroup symmetry. This idea was attacked by Rosenthal & Henderson [2003], arguing: *“However, a map with or without symmetry will be equally interpretable when the FSC is the same. Any threshold criterion that depends on the number of pixels in the map is not an absolute criterion for the evaluation of resolution*”. In reality, the model experiments in [van Heel & Schatz 2005] fully refutes their statement.

Böttcher B, Wynne SA, Crowther RA: **Determination of the fold of the core protein of hepatitis B virus by electron cryomicroscopy**. *Nature* 386 (1997) 88-91.

> This is one of the early cryo-EM papers in which a resolution better than 10Å was achieved for icosahedral particles. The FSC was used as a resolution metric in combination with an *ad-hoc* fixed 0.5 threshold criterion postulated without further statistical argumentation.

Conway J, Cheng N, Wingfield PT, Stahl SJ, Steven AC: **Visualisation of a 4-helix bundle in the hepatitis B virus capsid by cryo-electron microscopy**. *Nature* **386** (1997) 91–94.

> This is one of the early cryo-EM papers in which a resolution better than 10Å was achieved for icosahedral particles. The FSC was used as a resolution metric in combination with an *ad-hoc* fixed 0.5 threshold criterion postulated without further statistical argumentation.

Malhotra A, Penczek P, Agrawal RK, Gabashivili IS, Grassucci RA, Junemann R, Burkhardt N, Nierhaus KH, Frank J: **E. coli 70S ribosome at 15Å resolution by cryo-electron microscopy: Localization of fMet-tRNAfMet and fitting of L1 protein**. *J. Mol. Biol.* **280** (1998)103–115.

> With this paper the DPR [Frank 1981] has disappeared for this line of publications without discussion. This appears to be a tacit acceptance that the DPR, explicitly refuted in [van Heel 1987] was finally phased out in favour of the FRC/FSC. (However, the DPR issue was to resurface in [Liao 2010].) The FSC is introduced *without* any direct literature reference! The reference to the original FSC paper [Harauz & van Heel 1986] appears only a few lines later, and well as follows: “A *conservative approach (Böttcher et al., 1997; Conway et al., 1997) uses an FSC of 0.5 as the effective resolution of the reconstruction, while the three-sigma FSC criterion (Harauz & van Heel, 1986) uses a cutoff value that is close to zero in the interesting resolution range”.* This citation style is perplexing since neither the word “sigma” (nor symbol “σ”), nor the word “threshold” appear in the [Harauz & van Heel 1986] paper. Instead of scientifically defending their abandoning of the DPR metric, the authors move the discussion to the FSC *threshold* choice. They clearly prefer a “conservative” fixed 0.5 FSC threshold as per Penczek [1988], see below.

Penczek P: **Measures of resolution using Fourier shell correlation**. *J. Mol. Biol.* 280 (1998) 115–116. Appendix to: [Malhotra 1998]

> This paper attacks statements that nobody ever made, like: *“Sometimes the arguments for a cut-off equal to zero are put forward, supported by relatively weak statistical considerations*” [no references]. This argument is then literally taken over in their main paper [Malhotra 1998] for criticising the [Harauz & van Heel 1986] paper explicitly (see above). In attacking the non-existent proponents of that idea, the Frank & Al-Ali [1975] formula is referred to, without any restrictions attached to its validity. The author appears to consider the SNR (Signal to Noise Ration) the best of all possible metrics without any supporting evidence. The FSC is criticised by stating that it is difficult to convert to an SNR value using the Frank & Al-Ali [1975] formula, a formula that we have shown to be incorrect. An issue that is never mentioned in this paper is that the Frank & Al-Ali [1975] formula applies by their own statement to real-space images with **N** large (> 10.000) only. Because the relevant Fourier-space numbers of pixels/voxels ***N_r_*** (in the rings or shells) are so much smaller than the corresponding real space number **N**, the direct application of the Frank & Al-Ali formulas to the FRC/FSC is not justified.

Van Heel M: **Unveiling ribosomal structures: the final phases**. *Current Opinions in Structural Biology* **10** (2000) 259-264.

> With cryo-EM reaching resolution levels for ribosome structures better than ~10Å and X-ray crystallography achieving comparable resolution, a comparison between resolution metrics in both fields became opportune. It is argued that the FSC, which uses amplitudes and phases for comparing the data sets, is “a more direct measure of map quality” than the classical X-ray crystallographic R-factors.

Van Heel, M., Gowen, B., Matadeen, R., Orlova, E.V., Finn, R., Pape, T., Cohen, D., Stark, H., Schmidt, R., Schatz, M., and Patwardhan, A. **Single-particle cryo electron microscopy: towards atomic resolution**. *Quart. Rev. Biophys.* 2000b, **33**: 307-369.

> This review discusses the status of affairs in single-particle cryo EM at the turn of the century and gives a historical perspective on the resolution criteria in cryo-EM. To quote from this review: *“An interesting harmonisation has taken place in recent years in the use of resolution criteria for EM. After a long controversy, the differential phase residual DPR’ has been entirely superseded by the Fourier Shell Correlation criterion `FSC’”*
>
> A second quote: *“It unfortunately took some 15 years for the DPR’ school to (implicitly) admit its flaws by changing to the better FSC criterion*”

Grigorieff N, **Resolution measurement in structures derived from single particles** *Acta Cryst.* **D56** (2000) 1270-1277.

> This paper is primarily about the bias that the use of a common alignment reference can impose on a data set and not so much about resolution criteria themselves. The paper promotes the use of independent references for the alignment of single-particle half datasets in cryo-EM. This is thus one of the early papers promoting a “gold standard” independent alignment of half datasets. All resolution criteria can be used to illustrate this reference bias. “It is concluded that the alignment of images is always accompanied by a correlation of the noise and that this correlation is indistinguishable from a correlation arising from a signal”.

Frank J: **Single-particle imaging of macromolecules by cryo-electron microscopy** *Annu. Rev. Biophys. Biomol. Struct.* **31** (2002) 303–319.

> This review contains *“a note on resolution*” stating: “It is unfortunate, and quite confusing to non-specialists, that ***two different criteria of resolution*** are in use”. One would expect from this introductory sentence – and from the historical context – that the DPR [Frank 1981] would be compared with the FSC [Harauz 1986]. But actually this is a discussion about which FSC threshold to use: 0.5 [Frank 1988] or 3σ [Orlova 1997]. The DPR does not appear once in this review paper. This could be interpreted as a tacit admission that the DPR was poorly defined in [Frank 1981]. However, the DPR issue surfaced again in [Liao 2010].

Penczek PA: **Three-dimensional spectral signal-to-noise ratio for a class of reconstruction algorithms**. *J Struct. Biol.* **138** (2002) 34–46,

> The paper is an extension of the 2D SSNR [Unser 1987] to three dimensions. It therefore suffers from the same SSNR cross-terms problem as was discussed under [Unser 1987]. The Bershad & Rockmore [1974], and Frank & Al-Ali [1975] formula are also repeated explicitly here (formula “7”) without further discussion. This refutes the validity of the SSNR formulas derived in this paper.

Rosenthal PB, Henderson R: **Optimal determination of particle orientation, absolute hand, and contrast loss in single-particle electron cryomicroscopy**. *J. Mol. Biol.* **333** (2003) 721–745). *Appendix:* Rosenthal PB, Crowther RA, Henderson R: **An objective criterion for resolution assessment in single-particle electron Microscopy**. *J. Mol. Biol.* **333** (2003)743–745.

> The reasoning in this paper starts from the implicit assumption that only absolute, fixed-threshold value criteria can be correct and thus comes to statements of the nature: *“However, a map with or without symmetry will be equally interpretable when the FSC is the same. Any threshold criterion that depends on the number of pixels in the map is not an absolute criterion for the evaluation of resolution*”. The other basic assumption made is that the FOM used in X-ray crystallography is the standard by which to measure FSC results. In fact, the FSC is a better measure than are the traditional X-ray crystallography metrics as was discussed in the main paper (and in [van Heel 2000]). In deriving a significance threshold for the FSC, all cross-terms are explicitly assumed to be *orthogonal* (as in Bershad & Rockmore [1974]), thus removing all “√n” dependency from the results. The authors then proceed to postulate that *thus* all FSC threshold criteria should be fixed-value criteria: a dangerous circular argumentation. This paper is discussed in our main text.

Van Heel M., Schatz M: **Fourier Shell Correlation Threshold Criteria**, *J. Struct. Biol.* **151** (2005) 250-262.

> The Fourier Shell Correlation (FSC) criterion has by now become the accepted resolution function in cryo electron microscopy. Whereas σ-factor curves indicate the resolution level at which one has collected information significantly above the noise level, the ½-bit information curve indicates the resolution level at which *enough* information has been collected for a reliable interpretation. The important concept introduced here is the collecting of information measured in bits per voxel in Fourier space. As was already emphasized in this paper, the main problem with the literature is that the expectation values for the signal-versus-noise cross-terms are applied to the individual cross-terms. In this paper we speak of the “expected value” for the individual cross-terms. In two slips-of-the-tongue, the word “expectation” was written instead of “expected”. However, the model calculations in this paper unambiguously explicitly show what was meant in all cases. In this paper the errors in the literature, like in Frank & Al-Ali [1975] and Rosenthal & Henderson [2003], have been discussed extensively. Unfortunately, however, those insights have largely been ignored in the cryo-EM literature.

Baxter WT, Grassucci RA, Gao H, Frank J: **Determination of Signal-to-Noise Ratios and Spectral SNRs in cryo-EM low-dose imaging of molecules**. *J. Struct. Biol.* **166** (2009)126–132.

> This paper refers to the Frank & Al-Ali [1975] paper but no reference is made to the original Bershad & Rockmore [1974] work. The FRC/FSC metrics are mentioned without reference. The SSNR functions presented are derived from the FSCs through the incorrect Frank & Al-Ali [1975] formula in their Penczek [2002] form. All formulas in this paper ignore the cross-terms in the correlation function and are thus incorrect. The van Heel & Schatz [2005] paper, explicitly refuting the [1975] work, is ignored.

Liao HY, Frank J: **Definition and estimation of resolution in single-particle reconstructions**. *Structure* **18** (2010) 768-775.

> In this review paper, the authors for the first time acknowledge the problematic definition of the DPR [Frank 1981], thirty years earlier. Rather than accepting the corrected phase residual (FRPR, van Heel [1987], however, these authors choose to remedy the DPR flaws by an interesting construct: They suggest to loop over all possible linear scaling between the images and to then select the “best” phase residual value in their “*elastic-band”* approach to a reproducible DPR resolution metric. Note that it was already argued in [van Heel 1987], that even a corrected phase residual *(in casu,* the FRPR) is inferior to the FSC where both phases and amplitudes are taken into account.
>
> This paper also does not mention that the cited Frank & Al-Ali [1975] formulas were already refuted in [van Heel & Schatz, 2005]. The [2005] paper *is* referred to, but on an entirely different issue: “In contrast, in (van Heel and Schatz, 2005) it is *argued strongly in favour* of the criterion based on the 3σ.” This referencing is misleading, since in that [2005] paper the importance of the 3σ criterion was actually played down in favour of the ½-bit criterion.

Scheres SHW, Chen S: **Prevention of overfitting in cryo-EM structure determination**. *Nature Methods* **9** (2012) 853–854.

> This publication is discussed in our main paper: their *“gold standard*” reflects the cryo-EM specific data processing whereas the “FSC” referred to, is identically the same as the *“old standard*” FSC [Harauz & van Heel 1986], albeit used in connection with an incorrect 0.143 threshold. Note that the idea of performing independent “gold standard” alignments is much older, see for example [Grigorieff 2000] or [van Heel 2000b]. The “gold standard FSC” is a misleading nomenclature that confuses especially newcomers to the field.

Cardone G, Heymann JB, Steven AC. **One number does not fit all: mapping local variations in resolution in cryo-EM reconstructions, 184** (2013) 226–236.

> The authors apply a logical extension to the global FSC resolution metric by using the FSC locally to determine the resolution variations that occur locally within a larger 3D reconstruction. A fixed 0.5 FSC threshold is used, based on their considerations of the variance of the FSC metric. However, by using a *fixed* threshold value to compare the resolution of the sub-volumes with each other, the approach will necessarily yield inconsistent results. Performing FCSs on different subvolumes of a larger 3D reconstruction is like performing two independent FSC experiments. The metric used for such a comparison must be normalised correctly since a local FSC cannot even be compared to the global FSC of the same 3D reconstruction let alone to the FSC of a different sub-volume of the map. Fixed threshold values for the FSC are not properly normalised and thus cannot be used to compare the results of “different” FSC experiments. (The fixed 45° DPR threshold was criticized for the same reason [van Heel 1987]). Note that the [Cardone 2013] paper does not refer to the [van Heel & Schatz, 2005] paper specifying the ½ bit threshold curve.
>
> Comparability of global-volume FSC (360^3^ voxels, 0.6 Gaussian mask) with two different local-volume FSCs (24^3^ voxels, 0.6 Gaussian mask). The appropriate ½-bit threshold curves make the resolution values comparable, allowing for such metrics to be incorporated in automatic decision making such as needed for local resolution determination. All three curves show a high-resolution ½-bit cut-off at ~1/0.27=3.7Å, although the corresponding threshold levels differ from the full volume FSC to the local FSCs.
>
> 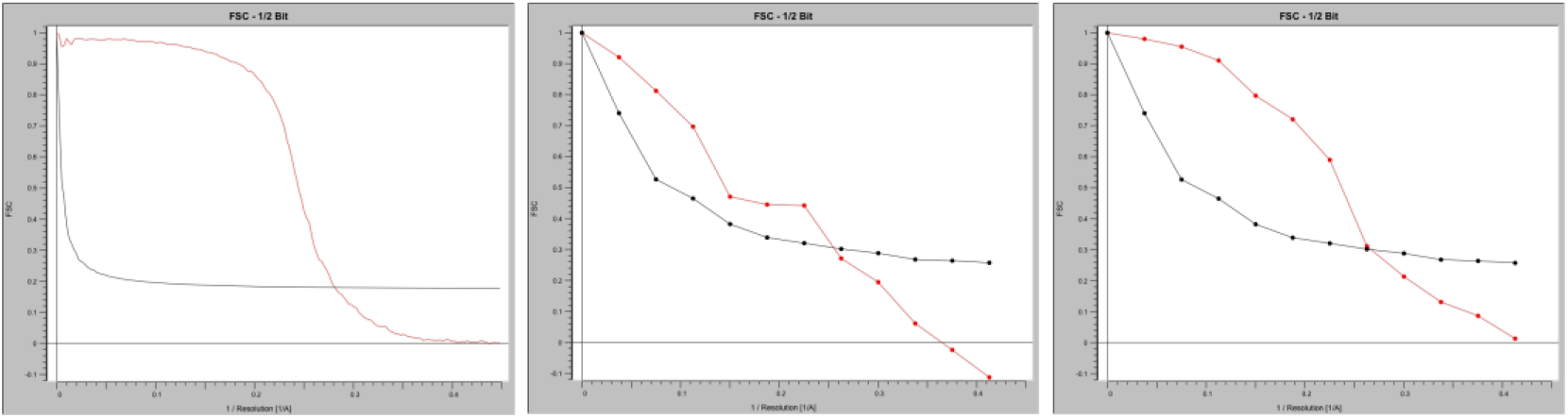

Chen S, McMullan G, Faruqi AR, Murshudov GN, Short JM, Scheres SHW, Henderson R: **High-resolution noise substitution to measure overfitting and validate resolution in 3D structure determination by single particle electron cryo-microscopy**. *Ultramicroscopy* **135** (2013) 24-35.

> The mathematics in this paper is again based on the assumed orthogonality of ***S***·*N*. Using the Bershad & Rockmore [1974] and Frank & Al-Ali [1975] arguments/formulas (without actually citing those papers) renders the resolution conclusions in this paper invalid. The basic idea of noise substitution is not invalidated but the paper needs to be fully rewritten based on a correct reproducible resolution metric.

Sorzano COS, Vargas J, Otón J, Abrishami V, de la Rosa-Trevín JM, Gómez-Blanco J, Vilas JL, Marabini R, Carazo JM: **A review of resolution measures and related aspects in 3D Electron Microscopy**. *Progress in Biophysics and Molecular Biology* **124** (2017) 1–30.

> A large number of metrics are discussed in this paper but no real conclusions are drawn. Virtually all formulas are based on expectation values applied to individual measurements, which, as we argued in the main paper, is the primary source of misunderstandings in cryo-EM quality metrics. The flawed Bershad & Rockmore [1974] and Frank & Al-Ali [1975] papers are cited throughout. The authors spend four pages of formulas on the DPR [Frank 1981], a metric that was explicitly refuted 30 years ago, and is no longer in use. The authors do cite the [van Heel & Schatz, 2005] paper and they criticize, that:
>
> *“one of the keystones of their* ([Van Heel & Schatz, 2005]) *reasoning is:*
>
>
> 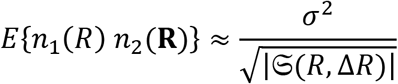
>
>
> *which is incorrect for independent random, zero-mean signals’*’
>
> In our [2005] paper we do have a similar formula (namely, formula “7”; originally due to Saxton [1978]) but that formula contains the *average* inner-product between two random vectors and no *expectation value* of that inner product:
>
>
> 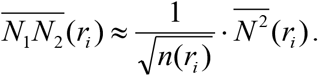
>
> There is a typo in our paper where the word “expected” has been switched by the word “expectation” in the text (not in formulas) but our formulas and model experiments leave no doubt as to what we are discussing. Sorzano *et al.* appear to come to conclusions that do not correlate with the arguments made in our paper.

Afanasyev P, Seer-Linnemayr C, Ravelli RBG, Matadeen R, De Carlo S, Alewijnse B, Portugal RV, Pannu NS, Schatz M, Van Heel M: **Single-particle cryo-EM using alignment by classification (ABC): Lumbricus terrestris hemoglobin at near-atomic resolution**, *IUCrJ* 4 (2017) 678-694. (<u>doi:10.1107/S2052252517010922</u>)

> Recent methodology developments in the IMAGIC-4D software system include an FSC as function of particle orientation. An improved resolution is observed in the preferred particle-orientation directions (and reduced one in less populated directions). One important aspect is the application of a correct threshold curve, i.e. the ½-bit curve. (All fixed-value threshold values are, again, inappropriate for directional FSCs). The figure (from the paper) includes the FSC_β<60°_ based on the preferred top-views (0°<β<60°); and an FSC_β>60°_ which assesses the contributions of the side-views (60°<β<90°). The circle indicates the area in which a high FSC value is a good indicator of the overall quality of the reconstruction.

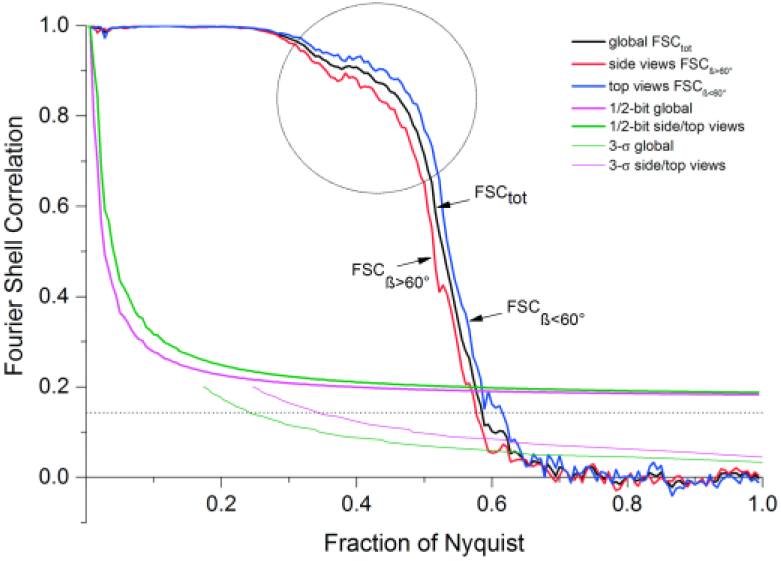

